## Supplementary Figures 1 -5 for "Thirteen independent genetic loci associated with preserved processing speed in a study of cognitive resilience in 330,097 individuals in the UK Biobank"

Table of contents

Supplementary Figure 1	GWAS outputs	2

Supplementary Figure 2	Heat map of genetic correlations	4

Supplementary Figure 3	Plot of 13 genetic loci	5

Supplementary Figure 4	Circos plots of associated chromosomes	12

Supplementary Figure 5	Gene mapping of RT and comparison to *Resilience*	17

**Supplementary Figure 1:** Manhattan plots for (a) EY+Res, (b) EY/NonRes, (c) EY, (d) RT (n=164,000) (e) RT (n =333,664) (f) EduYears (g) Manhattan plot and QQ plot for cognitive change in the Health and Retirement Study.

EY+Res and EY/NonRes were the input GWASs for GWAS-By-Subtraction (GBS) that generated the *Resilience* GWAS and the *EduYears* GWAS.

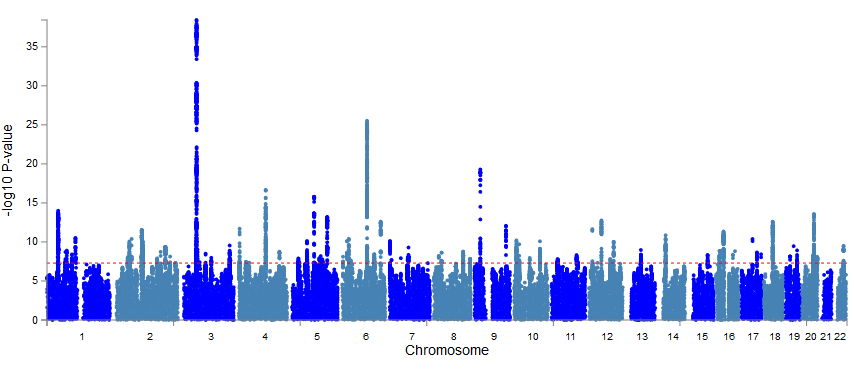

a. EY+Res (n = 156,011)

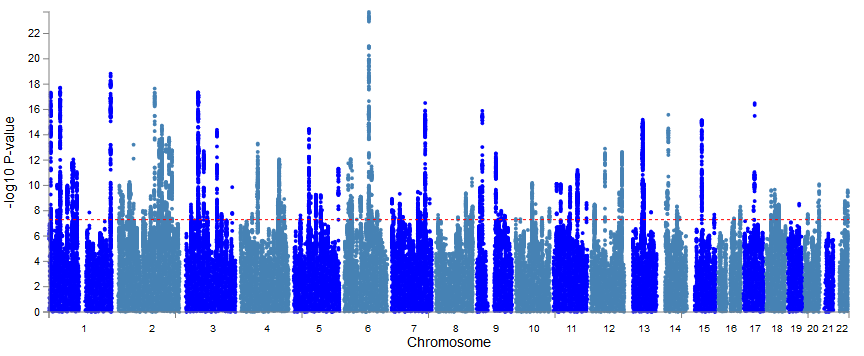

b. EY/NonRes (n = 174,086)

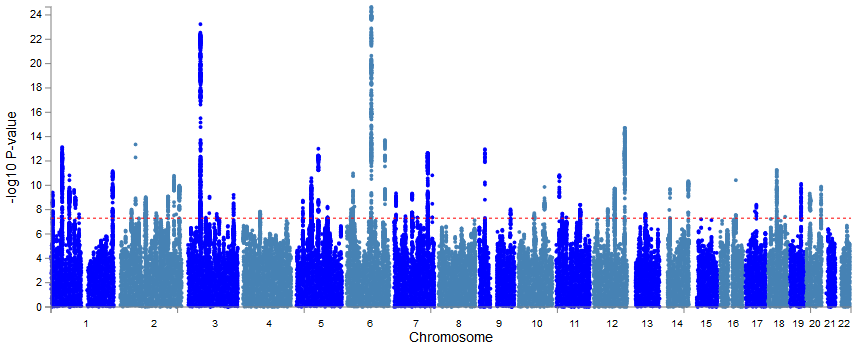
c. EY (n = 164,000)

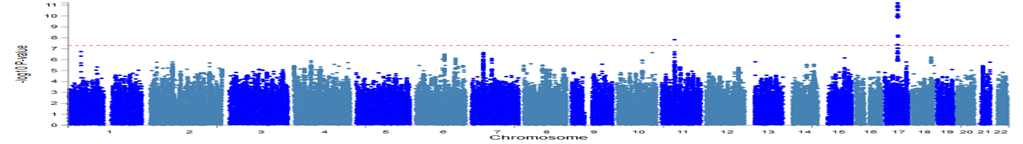

d. RT (n = 164,000)
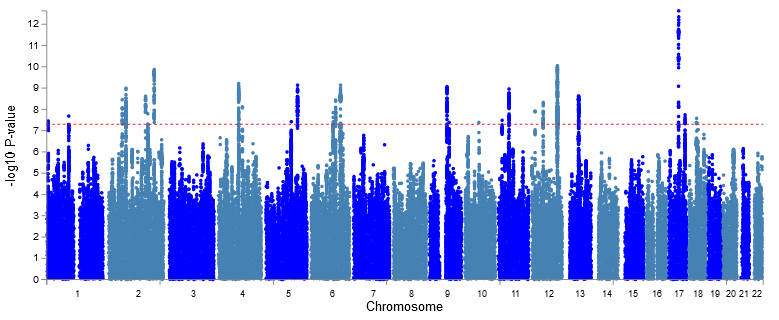

e. RT (n = 333,664)

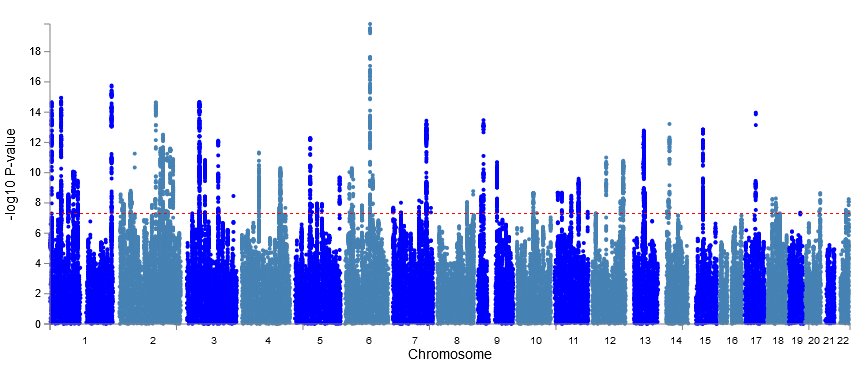

f. *EduYears* (n = 166,122)

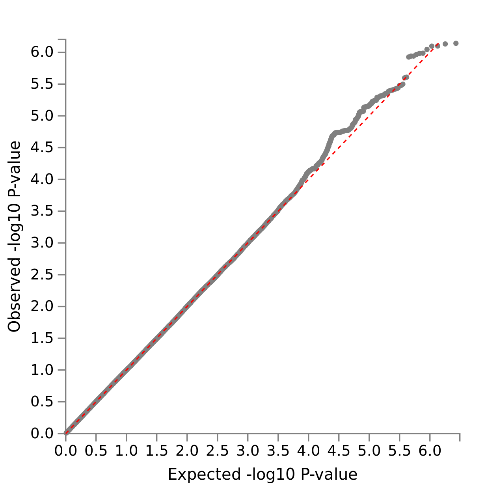

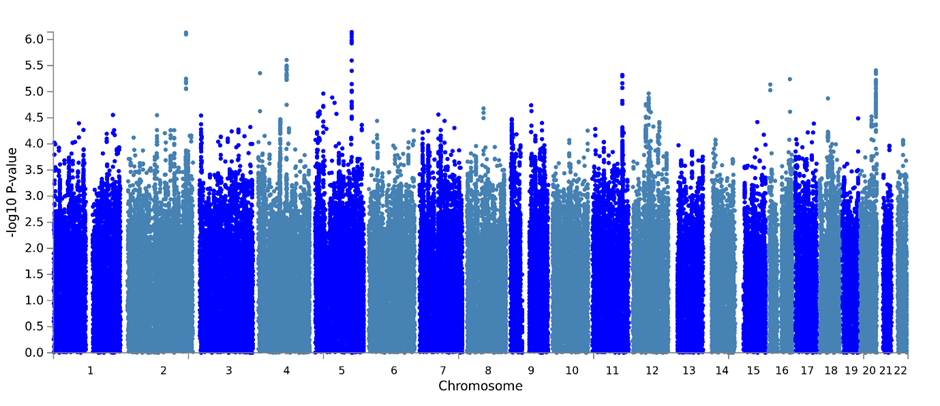

g. QQ plot and manhattan plot of cognitive change in the Health and Retirement Study.

**Supplementary Figure 2:** *Heat map showing genetic correlations between the two GBS GWAS of Resilience and EduYears, the two inputs to GBS (EY+Res and EY/NonRes) and GWAS of the two variables used to create these phenotypes (EY and RT).*

|  | *Resilience* | *EduYears* | EY+Res | EY/NonRes | EY | RT |  |
| --- | --- | --- | --- | --- | --- | --- | --- |
| *Resilience* |  |  |  |  |  |  |  |
| *EduYears* | 0.01* |  |  |  |  |  | 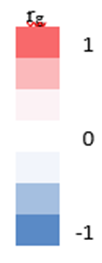 |
| EY+Res | 0.84 *** | 0.55*** |  |  |  |  |  |
| EY/NonRes | 0.01* | 1.0*** | 0.54*** |  |  |  |  |
| EY | -0.47*** | -0.9*** | -0.88*** | -0.89*** |  |  |  |
| RT | 0.8*** | -0.56*** | 0.36*** | -0.56*** | 0.14** |  |  |

*** P<1x 10 ^-10^, **P< .0002, * Not significant.

**Supplementary Figure 3:** Plots of the 13 independent genetic loci (a - m) showing the top lead SNPs, lead SNPs, independent significant SNPs and associated genes.

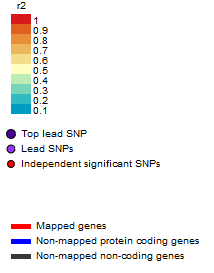

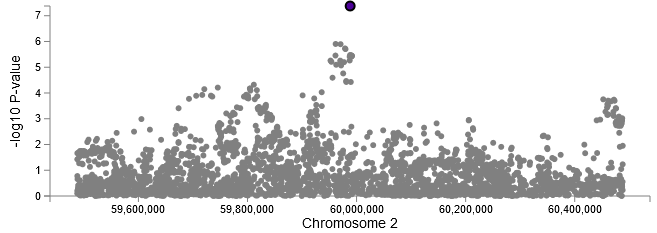

Locus 2: Chr2:59987310-59988258

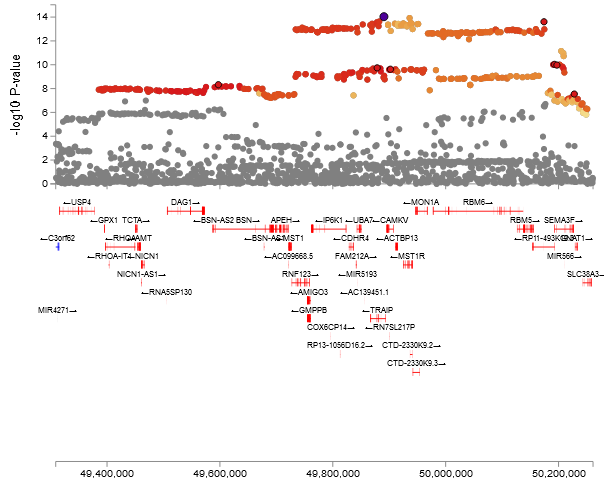

a. Locus 3: Chr3:49385417-50248954

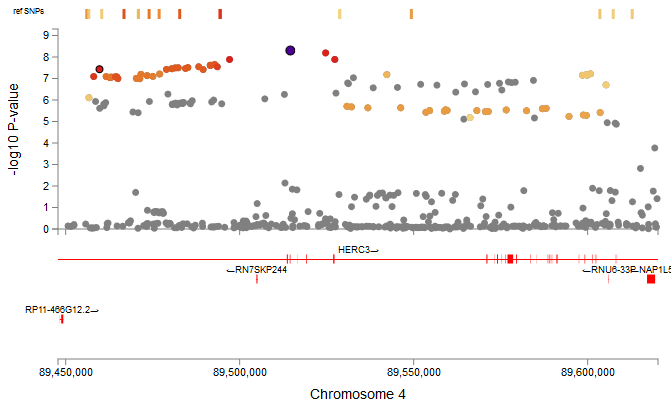

b. Locus 4A: Chr4:89455635-89612380

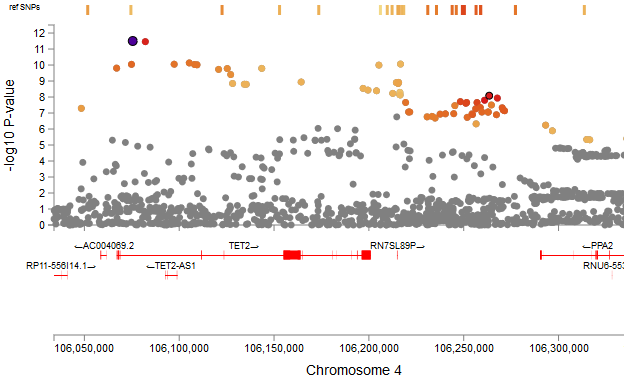

c. Locus 4B: Chr4:106048360-106335951

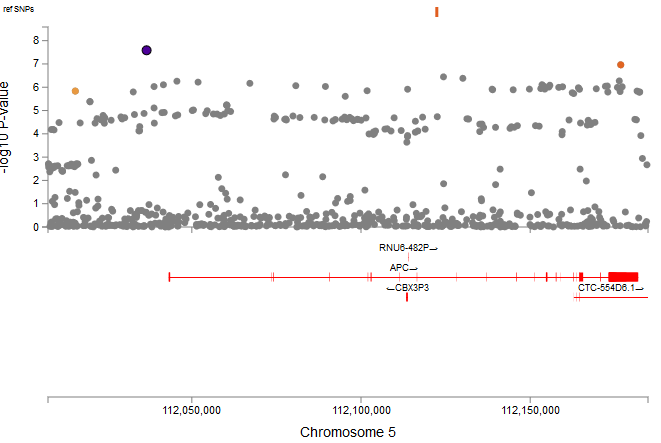

d. Locus 5A Chr5:112015555-112176756

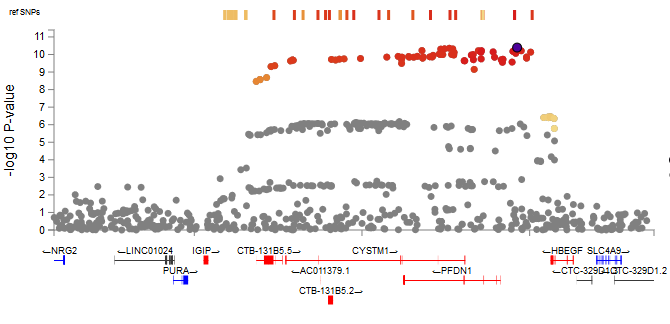

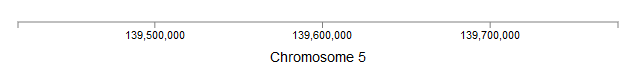

e. Locus 5B: Chr5:139517197-139714690

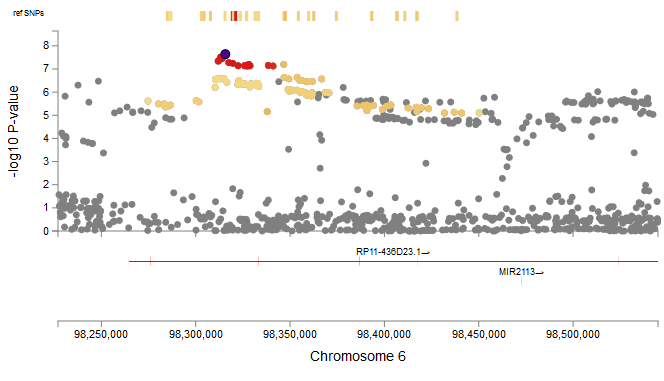

f. Locus 6: Chr6:98274701-98450190

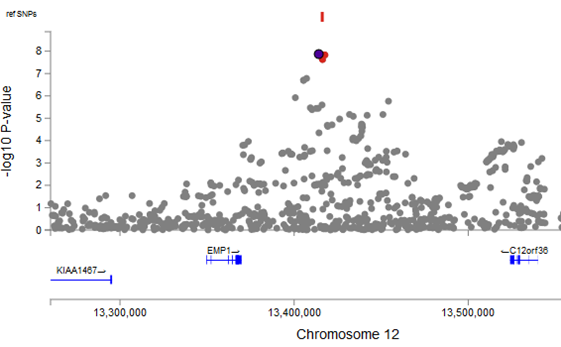

g. Locus 12A: Chr12:13414139-13417617

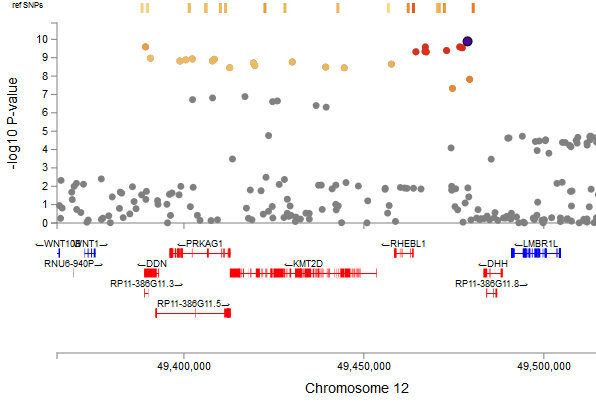

h. Locus 12B: Chr12:49387955-49479968

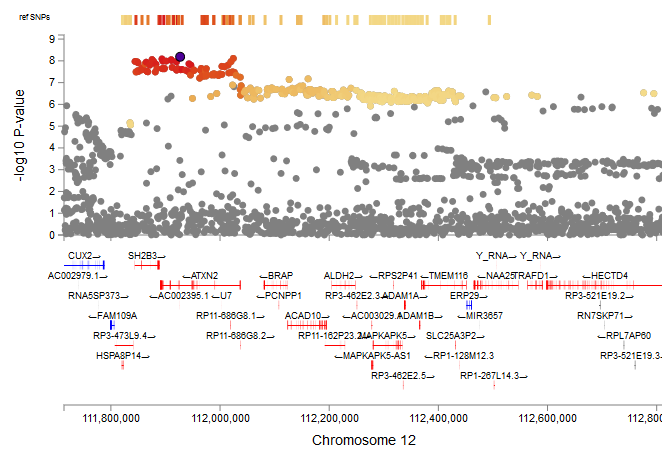

i. Locus 12C: Chr12:111818487-112817847

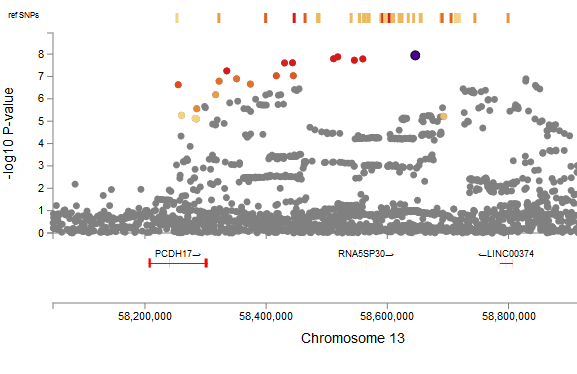

j. Locus 13: Chr13:58250322-58796832

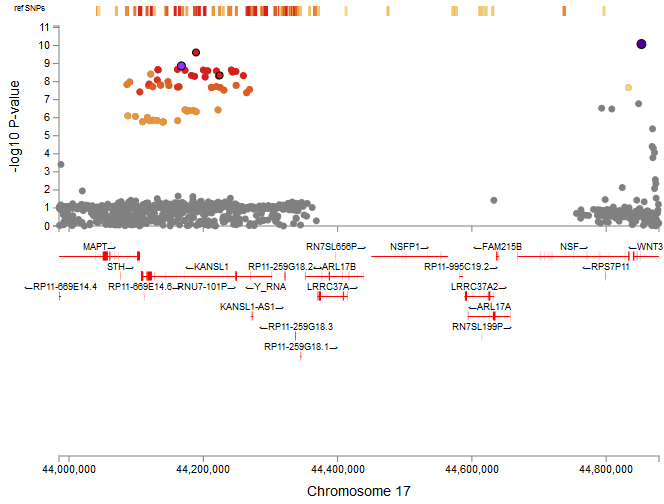

k. Locus 17: Chr17:44040184-44852612

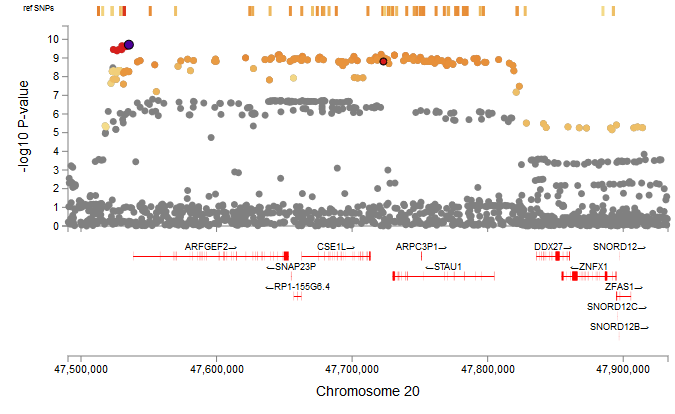

l. Locus 20: Chr20:47511792-47914180

**Supplementary Figure 4**: Circos plots of chromosomes that contain genome-wide significant loci. Genomic loci are highlighted in blue. Orange represents genes that are mapped by chromatin interaction and green represents eQTL mapping. If genes are mapped by both, they are highlighted in red. The dark blue portion of the inner circles represents the loci.

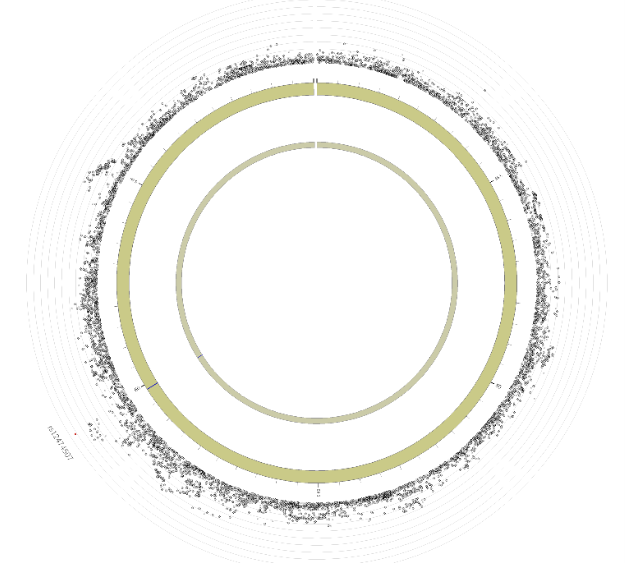

Chromosome 2

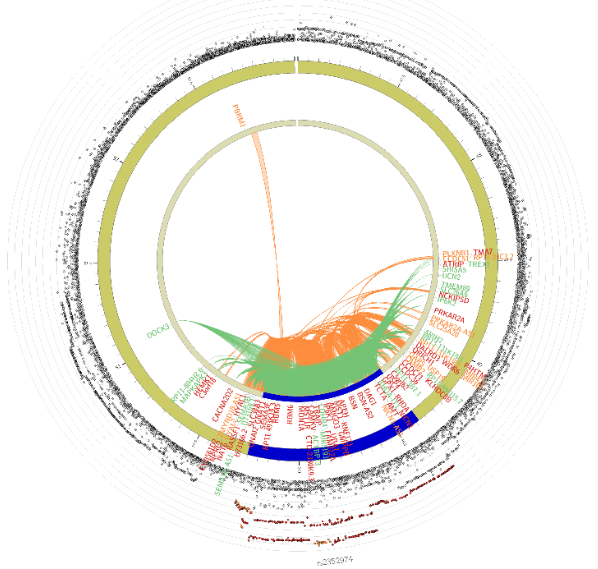

Chromosome 3

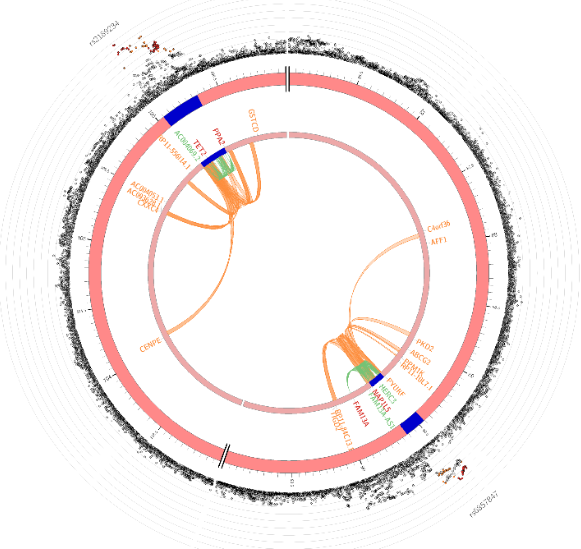

Chromosome 4

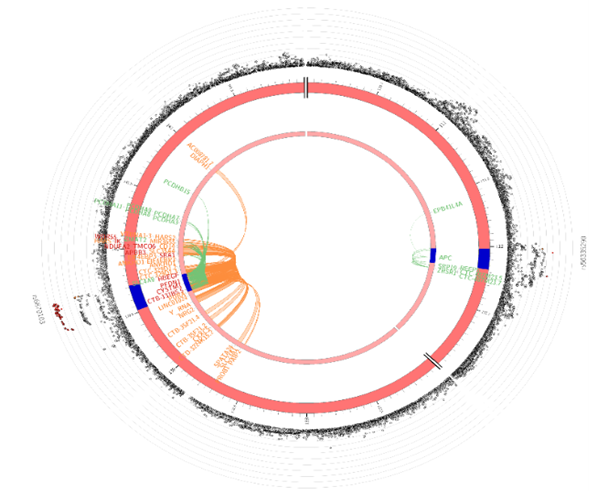

Chromosome 5

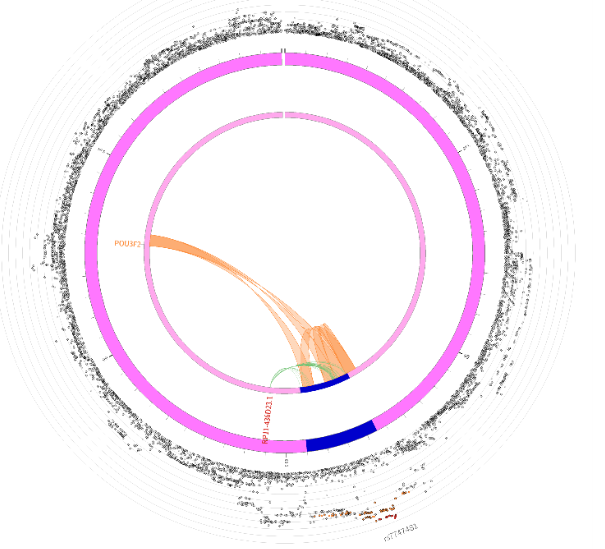

Chromosome 6

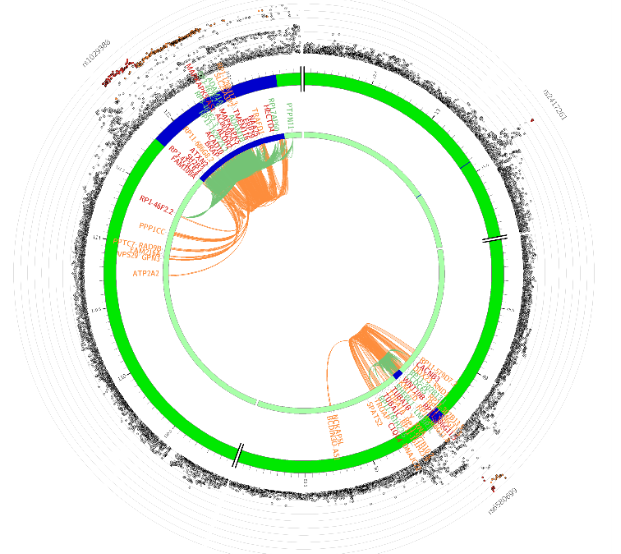

Chromosome 12

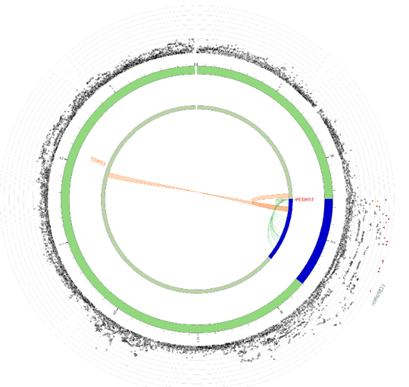

Chromosome 13

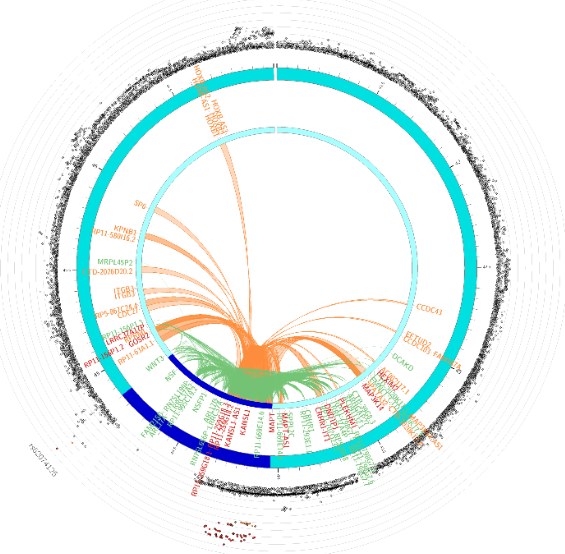

Chromosome 17

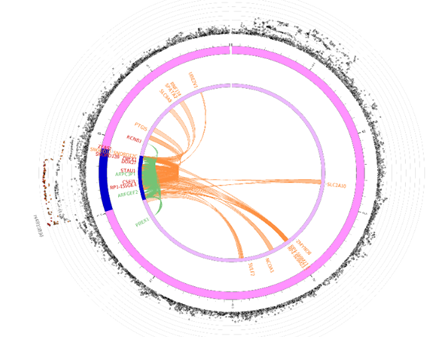

Chromosome 20

**Supplementary figure 5:** Gene mapping of RT and comparison to *Resilience.* **a** Venn diagram of overlapping mapped genes by four strategies showing 27 genes were mapped by all four strategies for RT. **b** Venn diagram showing the overlap between the 27 prioritized genes from functional analysis of RT and the 33 prioritized genes from functional analysis of *Resilience.*

**a**

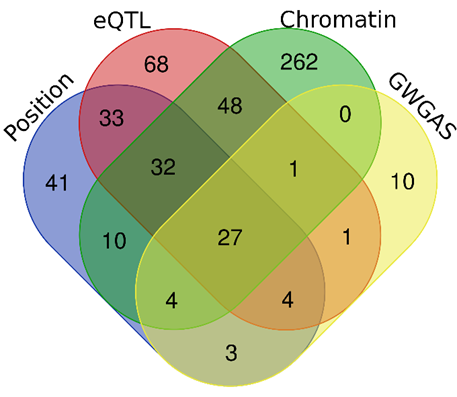

**b**

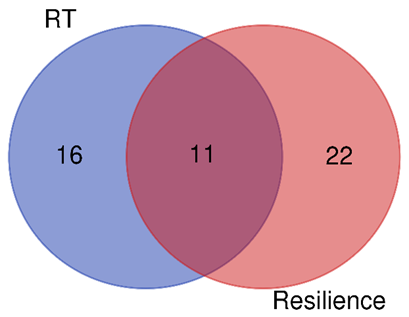
